# Analysis of Mouse Vocal Communication (AMVOC): A deep, unsupervised method for rapid detection, analysis, and classification of ultrasonic vocalizations

**DOI:** 10.1101/2021.08.13.456283

**Authors:** Vasiliki Stoumpou, César D. M. Vargas, Peter F. Schade, Theodoros Giannakopoulos, Erich D. Jarvis

## Abstract

Some aspects of the neural mechanisms underlying mouse ultrasonic vocalizations (USVs) are a useful model for the neurobiology of human speech and speech-related disorders. Much of the research on vocalizations and USVs is limited to offline methods and supervised classification of USVs, hindering the discovery of new types of vocalizations and the study of real-time free behavior. To address these issues, we developed AMVOC (Analysis of Mouse VOcal Communication) as a free, open-source software to analyze and detect USVs in both online and offline modes. When compared to hand-annotated ground-truth USV data, AMVOC’s detection functionality (both offline and online) has high accuracy, and outperforms leading methods in noisy conditions, thus allowing for broader experimental use. AMVOC also includes the implementation of an unsupervised deep learning approach that facilitates discovery and analysis of USV data by clustering USVs using latent features extracted by a convolutional autoencoder and isimplemented in a graphical user interface (GUI), also enabling user’s evaluation. These results can be used to explore the vocal repertoire space of the analyzed vocalizations. In this way, AMVOC will facilitate vocal analyses in a broader range of experimental conditions and allow users to develop previously inaccessible experimental designs for the study of mouse vocal behavior.

## Introduction

Over the past two decades there has been a growing interest in the usage and signaling of vocalizations in mice, with large efforts going into studying the underlying neurobiological mechanisms for auditory processing (***Pomerantz et al., 1983***; ***Liu et al., 2003***; ***Neilans et al., 2014***; ***Perrodin et al., 2020***; ***Holy and Guo, 2005***), and the production of vocalizations (***Arriaga et al., 2012***; ***Chabout et al., 2016***; ***Okobi et al., 2019***; ***Zimmer et al., 2019***; ***Gao et al., 2019***; ***Tschida et al., 2019***; ***Michael et al., 2020***). The tools available for experiments in mice provide a promising model for studying the neural basis of vocalizations, as well as the effects of genes on the origin and development of vocal and neural anatomy (***Grimsley et al., 2011***; ***Bowers et al., 2013***; ***Chabout et al., 2016***; ***Tabler et al., 2017***).

Mice produce ultrasonic vocalizations (USVs; 30-110kHz) relative to the human hearing range (2-20kHz). Both sexes of mice produce USVs from an early neonatal age through adulthood (***Grimsley et al., 2011***). Efforts have been made to better understand these USVs in terms of their structure, classifying their multi-syllabic structure, as well as the non-random sequencing of syllables (***Holy and Guo, 2005***; ***Chabout et al., 2016***; ***Calbick et al., 2017***; ***Castellucci et al., 2018***; ***Hertz et al., 2020***). USVs exhibit considerable variation across inbred strains such that innate USV repertoires can be used as a phenotyping marker for different genotypes (***Melotti et al., 2021***), and as a behavioral readout for genetic modifications and speech related mutations (***Scattoni et al., 2009***; ***Chabout et al., 2016***; ***Castellucci et al., 2016***). USVs can also vary within individuals based on social environment and affective state. For example, adult male mice are known to modify the temporal organization and the rate of vocalizations for different social contexts (***Chabout et al., 2015***). While we have learned a lot from these behavioral and genetic studies, we still do not know very much about the ways in which the murine brain generates and processes the content and variation of conspecific USVs. For these reasons, in order to advance our understanding of the neurobiology of vocalizations, it is equally important to more precisely understand the vocalizations themselves as units of behavior.

Historically, most vocal research was done by hand-annotating spectrograms. Successive technological improvements have led to many new automated methods that allow for unsupervised detection and classification of vocalizations.

These advances in vocal detection include tools developed specifically for analyzing the USV repertoire of mice. Early tools used supervised classification methods that rely explicitly on the acoustic parameters of the recordings. One such tool is Mouse Song Analyzer (MSA) (***Holy and Guo, 2005***; ***Arriaga et al., 2012***; ***Chabout et al., 2015***). MSA first generates a spectrogram from the audio recording, that is subsequently thresholded to remove white noise. Frequencies outside of mouse USV range of 35-125 kHz are discarded. The spectrogram is used to compute frequency, amplitude, spectral purity, and discontinuity across the entire recording. A combination of user-defined thresholds for each of these features is used to detect USVs. Lastly a duration filter is applied to remove any detected sounds that fall below a predefined value, all remaining detections are considered USVs. The detected USVs are then classified based on the number of gaps, or “pitch jumps”, that are present within the detected USVs (***Chabout et al., 2015***). USVSEG (***Tachibana et al.*** (***2020***)) is also a MATLAB tool used for vocalization detection, emphasizing on noise removal of the spectrogram in the cepstral domain before thresholding to detect whether a segment contains a vocalization or not. Another tool is Ax (***Neunuebel et al.*** (***2015***)), which seeks to detect vocal signals by keeping time-frequency points of the spectrogram that signifcantly exceed noise values.

As new methods have arisen for the unsupervised assessment of USVs (***Van Segbroeck et al., 2017***), some have implemented machine learning and neural networks in their processes (***Coffey et al., 2019***; ***Fonseca et al., 2021***). Machine learning methods have been used improving detection accuracy, but performance is limited by how well a network is trained, and it may not always generalize across experiments.

This includes Mouse Ultrasonic Profile ExTraction (MUPET), a MATLAB open source tool, developed by ***Van Segbroeck et al.*** (***2017***) to detect syllables, analyze the vocalizations features and cluster the syllables depending on these features. MUPET first filters the signal to keep high frequencies (25-125 kHz). It then uses spectral subtraction to remove stationary noise, and at last it computes the power of the spectral energy in the ultrasonic range above a specific threshold. The vocalizations are converted to representations by using negative matrix factorization (NMF) and gammatone filters. The filtered spectrograms are then used to cluster vocalizations based on spectral shape similarities. The clustering is done by using K-Means, and user-defined number of “repertoire units” (clusters). The authors note that MUPET can also be used with many non-rodent species’ vocalizations.

DeepSqueak is a software suite for USVs detection and analysis (***Coffey et al.*** (***2019***)). It splits the recording into areas of interest, computes the corresponding sonograms and passes them to a Faster-RCNN (recurrent convolutional Neural Network) object detector, which consists of two networks. The first network is a region detection network, which proposes sections of the spectrogram that could contain actual vocalizations. These sections are then used as inputs to a second network, a convolutional neural network (CNN), and are classified depending on whether or not the sections contain vocalizations. This process has been recently updated to use a You Only Look Once (YOLO) network to improve detection quality. These networks can also be trained on new vocalizations. DeepSqueak can also be used for clustering the detected syllables, either with a supervised or unsupervised method. Their unsupervised approach gives the user the opportunity to define three weighted input features: shape, frequency and duration of the vocalization. An important difference from MUPET is that the clustering function of MUPET takes syllable amplitude into account, whereas DeepSqueak does not, which can be considered an advantage for DeepSqueak, given that the amplitude of a vocalization can depend on the recording setup itself, among other factors.

VocalMat is a MATLAB tool (***Fonseca et al.*** (***2021***)), which uses image processing techniques and differential geometry analysis on the spectrogram of a recording to detect vocalization candidates. It can then classify detected USVs into 12 predefined categories (including noise), by using a CNN. As with DeepSqueak, VocalMat’s networks can be retrained as well.

A more recent tool is Deep Song Segmenter (DeepSS), which has been used for annotation of songs of mice, birds and flies (***Steinfath et al., 2021***). DeepSS learns a representation of sounds features directly from raw audio recordings using temporal convolutional networks (TCNs), based on dilated convolutions. It is a comparatively fast, supervised annotation method, since the network is trained with manually annotated recordings. It can also be combined with unsupervised approaches to reduce the amount of manual annotation required.

There are, however, limitations to each of these approaches. The supervised classification methods, like MSA (***Holy and Guo, 2005***; ***Arriaga et al., 2012***; ***Chabout et al., 2015***), limits their utility in being able to analyze datasets for changes or novelty in the USV repertoire across individuals and genotypes. All methods thus far do not perform analyses in real time. To address these pitfalls, we sought to create a tool for mouse USVs that is both computationally efficient and accurate, and that can provide a less biased classification of USVs.

In this work we present Analysis of Mouse Vocal Communication (AMVOC), a new open-source tool for mouse USV research. AMVOC’s purpose is twofold: (a) it uses dynamic spectral thresholding to *detect* the presence of USVs in audio recordings (both offline and online), and (b) similar to other machine learning approaches it analyzes and visualizes the detected USVs using feature representations extracted from deep convolutional autoencoders, and uses these feature representations for *clustering* and data exploration. AMVOC’s ability to detect USVs has been extensively evaluated using real recordings, and AMVOC’s accuracy has been shown to outperform most state-of-the-art tools in various acoustic environments, allowing for more flexible experimental set ups. In addition, while many of the other USV detection tools available are specifically developed for offline analysis of vocalizations, AMVOC is unique in that it can also be used to detect and measure USVs in real-time with an accuracy that rivals the accuracy of offline approaches and detection speeds at behaviorally relevant timescales. Lastly, the proposed deep feature extraction technique, using a convolutional autoencoder, produces unsupervised USV classifications that can be used as a basis to discover biologically relevant USV clusters. This provides an unparalleled usage that opens up new avenues to better understand mouse vocal behavior and its associated neurobiology.

## Results

### Experimental evaluation of the AMVOC detection method

Our objective was to design and implement a robust (in terms of detection performance) but also computationally efficient USV detection method, as our vision was to build a real-time, online pipeline. Because of the online vision of the design, we wanted to ensure that our method could still perform with high accuracy despite the demands of online processing.

We compared our proposed detection methodology with several popular USV detection tools, such as MUPET (***Van Segbroeck et al., 2017***), VocalMat (***Fonseca et al., 2021***), and DeepSqueak (***Coffey et al., 2019***). We also wanted to compare our new tool with MSA, which was developed in our lab. We rewrote MSA, from the original MATLAB implementation (MSA1) to a Python implementation (MSA2) with added filtering components to improve USV detection rates. We noted that MSA1 was cutting off parts of beginning of syllables, underrepresenting the full spectral duration of the USV syllables, as well as missrepresenting the timestamps of the detected USVs. MSA2 first bandpasses the raw audio (30-115 KHz) and then generates a spectrogram. The spectrogram is thresholded according to the signal-to-noise ratio at each frequency.

Due to the vast range of possible parameters that could be tuned in each method we compared against, we used default settings, unless there were other documented settings that were used (***Chabout et al., 2017***). In order to evaluate and compare the aforementioned methods, we used Dataset D1, which consists of 9 audio segments of 5-10 seconds each, containing 245 annotated syllables in total from 14 mice (see Section).

To evaluate the range of experimental contexts recordings could be taken from, we split the recordings into two categories: normal and noisy. Normal parts of the recording are the ones where the vocalization detection is relatively straightforward, because the energy easily surpasses the background energy. Noisy parts contain background noise, such as cage bedding or physical interacitons between the mice, any of which makes the detection more difficult and ambiguous, even for the human eye observing the raw spectrograms. The evaluation metrics are calculated separately for the two categories.

For evaluation and comparison we adopted two performance metrics, namely the temporal F1 and the event F1 score:

- For the *temporal F1 score*, we interpret the vocalization detection as a classification task of each 1 ms time frame into two categories (vocalization or no vocalization). Then we calculate the precision and recall rates by comparing the detected vocalizations to the ground-truth vocalizations. Their harmonic mean is the temporal F1 score.
- The *event F1 score* is the harmonic mean of the two following fractions: (a) the number of events a method detected that are annotated in the ground-truth data by the number of events detected by a method (i.e. precision), and (b) the number of events annotated in ground-truth data that are detected with a method by the number of events of ground-truth annotations (i.e. recall).

Using Dataset D1, we found that AMVOC outperforms the other methods with respect to event F1 score, both in clean and noisy segments of the recordings, whereas MSA2 and DeepSqueak performed slightly better than the others with respect to temporal F1 score (Table 1), largely due to the more successful detection in the noisy parts of the recordings.

**Table 1.** F1 scores of our proposed method and other methods.

| F1 score |  | AMVOC |  |  |  |  |  |  |
| --- | --- | --- | --- | --- | --- | --- | --- | --- |
|  |  | offline | online | MSA1 | MSA2 | MUPET | VocalMat | DeepSqueak |
| Temporal | Normal | 84 | 85 | 46 | 88 | 85 | **91** | 83 |
|  | Noisy | 67 | 68 | 23 | 71 | 53 | 57 | **76** |
|  | **Average** | 75.5 | 76.5 | 34.5 | **79.5** | 69 | 74 | **79.5** |
| Event | Normal | **97** | **97** | 66 | 94 | 93 | 90 | 93 |
|  | Noisy | **84** | 83 | 33 | 72 | 57 | 58 | 81 |
|  | **Average** | **90.5** | 90 | 49.5 | 83 | 75 | 74 | 87 |

We also assessed the trade-off between processing speed and detection by determining the processing ratio for AMVOC and each method we compared to (Table 2). This metric provides an assessment of how quickly a method is able to process a batch of audio data, which is important for considering the feasibility of real-time processing. AMVOC had an intermediate real-time processing ratio to detect the vocalizations (Table 2). MUPET was the fastest method, whereas VocalMat and DeepSqueak were the slowest (Table 2). The reason for the latter two methods being slower is likely due to their image processing steps used to detect USVs. It is also meaningful that we take into account both of the two aforementioned metrics, since it may be important for particular experimental requirements that a certain method combines accurate and fast detection. We compared the average F1 score with the real-time Processing Ratio for both temporal and event F1 scores (Figure 1A and B). AMVOC and DeepSqueak achieved the highest temporal and event F1 score, but AMVOC had a considerably better time performance relative to DeepSqueak. While both AMVOC and DeepSqueak were the most accurate overall, AMVOC performed better in segments where noise energy was near that of the USVs (Figure 1C).

**Table 2.** Real-time Processing Ratio of all compared methods. Real-time processing ratio is defined as 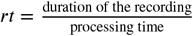 and is shown for each method. The processing time is calculated as the time needed to just detect the USVs. The experiments carried out to compute the real-time ratio were executed for 5 different recordings, 3 times for each, and the average time for each method was calculated. Obviously, a high real-time processing ratio means that a small processing time is required in order to detect the vocalizations of a certain signal (e.g. *rt* = 30 means that the respective method is 30 times faster than real-time, meaning it takes 1 minute to process 30 minutes of audio information).

| Methods | Real-time Processing Ratio |
| --- | --- |
| MUPET | **32.4** |
| MSA2 | 29.9 |
| MSA1 | 28.1 |
| AMVOC | 21.2 |
| DeepSqueak | 8.2 |
| VocalMat | 4.3 |

**Figure 1.**
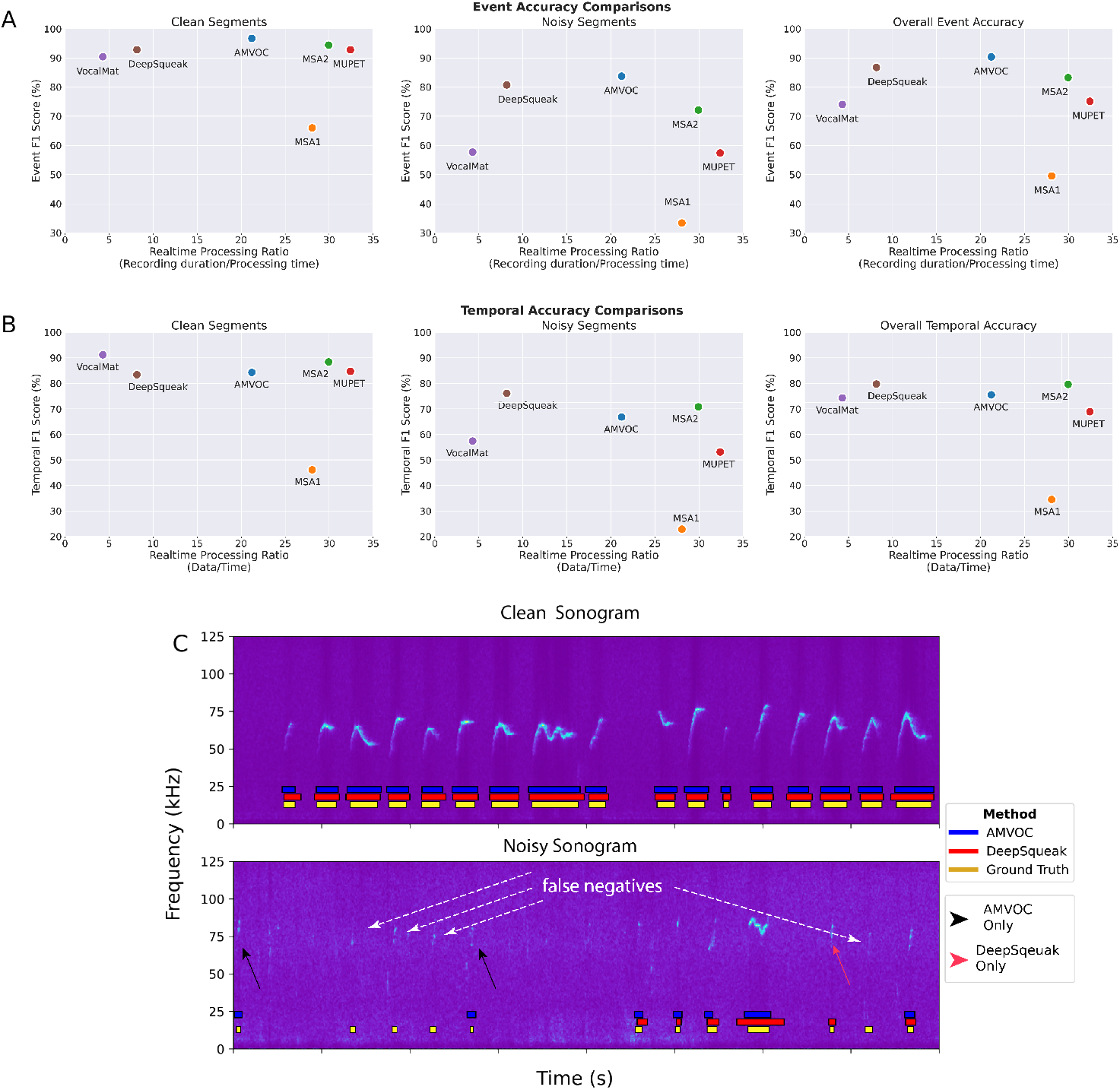
Accuracy of AMVOC and Other Methods A-B) Event and temporal accuracy of different USV detection methods compared against our ground truth data in different qualities of recordings. C) Examples of the two highest performing methods, AMVOC and Deepsqueak, on clean (top) and noisy (bottom) segments of USV recordings. Contrast modified to facilitate visualization in the noisy spectrogram.

To mitigate bias in our evaluation of AMVOC, we also compared AMVOC with the Dataset published alongside VocalMat (hereafter referred to as VM1) by ***Fonseca et al.*** (***2021***). VM1 consists of 7 different recordings from 7 mice (5-15 days old, of both sexes) (***Fonseca et al., 2021***). We did not change AMVOC’s pre-determined configuration (parameters *t* and *f*) for this evaluation, in order to examine how robust the selection of *t* and *f* is for recordings produced in different conditions.

The recordings of VM1 contain a constant noise artifact at 30 kHz, to remove unwanted distortions we used a high pass filter with a higher cut-off frequency than before (45 kHz). By implementing a 45 kHz cut-off, the new range we examined for detection in VM1 was between 45 and 110 kHz, instead of 30-110 kHz. Because the ground truth annotations by ***Fonseca et al.*** (***2021***) only declare the start time of each vocalization, we examined start times of the different methods compared to ground truth start times with 20 ms tolerance, similar to what Fonseca et al. used in their assessment of VocalMat vs DeepSqueak. We also considered a ground truth vocalization as “found” when the ground truth start time was between the start and end time of a detected vocalization (in case some method merged successive vocalizations). Since we only have ground truth start times, we only used event and not temporal evaluation. We calculated precision, recall and F1-score of VM1 detection results as we had done with Dataset D1, for each of the seven recordings separately, and then their mean.

The detected vocalizations of VocalMat are taken from ***Fonseca et al.*** (***2021***), whereas we ran the experiments with MUPET, DeepSqueak and, of course, AMVOC.

Without tuning on this specific dataset, AMVOC (both offline *and* online) achieved the best results compared to the other methods except for VocalMat (Table 3). Notably, VocalMat’s detection method is trained on data similar to test data from Dataset VM1, so high detection quality by VocalMat on their own data is expected.

**Table 3.** Event F1-score of all methods. All measurments provided as mean and standard error of the mean (SEM) from *n* = 7 samples (recordings).

| Methods | Event F1-score (%) |
| --- | --- |
| AMVOC offline | 95.6 (+/- 2.5) |
| AMVOC online | 93.6 (+/- 3.8) |
| VocalMat | **98.5 (+/-0.6)** |
| MUPET | 76.0 (+/-10.2) |
| DeepSqueak | 85.1 (+/-3.8) |

### Experimental evaluation of the AMVOC clustering method

After evaluating the segmentation and detection performance of AMVOC we wanted to develop a way to assess the repertoire composition of the USVs. To do so, we developed an autoencoder approach (see Section) whereby latent features are extracted from images of detected USVs, transformed, and then clustered for manual inspection and evaluation (Figure 2). Although others have shown autoencoders can extract relevant information about an individual mouse’s vocalizations (***Goffinet et al., 2021***), similarly, we wanted to ensure that the autoencoder we designed could capture features of USVs that may not be obviously calculated or discerned relative to what could be obtained from simple feature descriptions that are often used for comparing USVs of different mice.

**Figure 2.**
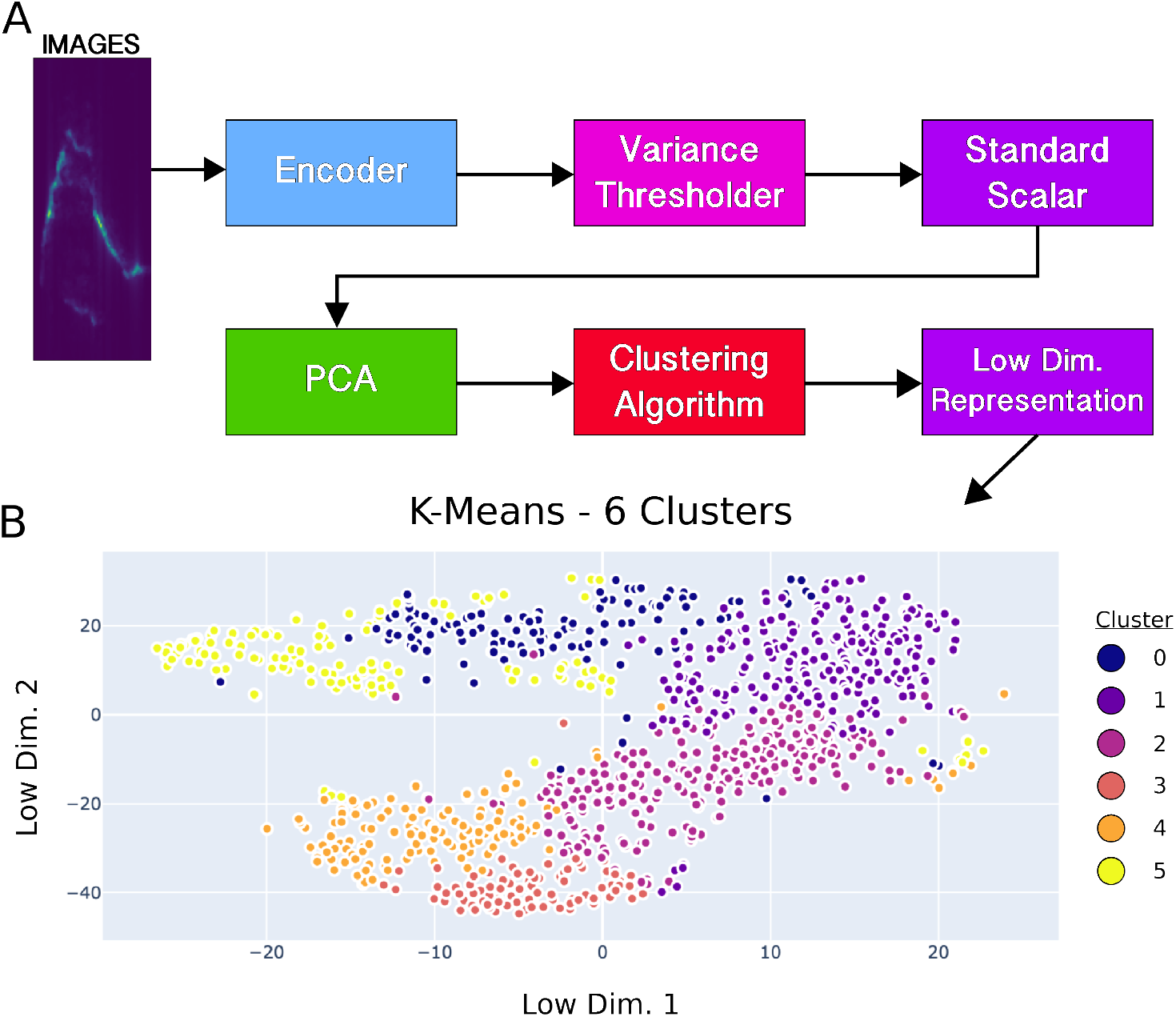
Overview of deep feature extraction procedure. A) Flow diagram of the general procedures used to take image data from USV spectrograms into clusters. B) Cluster example using deep features with K-Means clustering and 6 clusters.

Our primary goal is to compare the deep feature extraction method (see Section) to the baseline feature extraction method using hand-picked features (e.g. bandwidth, duration, frequency max, and frequency min) (described in methods Section). We compared these two feature extraction approaches by evaluating the derived clustering of the two kinds of features. We evaluated how well the different clustering methods perform in grouping vocalizations with common features. To achieve these two goals, we used data from four different annotators, two of whom are domain experts, and the other two are not. By using the AMVOC GUI, each annotator evaluated the 4 recordings from Dataset D3, which were generated by selecting segments of 72 different recordings (). The annotators evaluated 3 different clustering configurations for each recording: Agglomerative clustering with 6 clusters, Gaussian Mixture clustering with 6 clusters and K-Means clustering with 6 clusters.

In order to ensure impartiality and objectivity, the annotators evaluated the clustering derived from both feature extraction methods (named as Method 1 for deep features and Method 2 for hand-picked features), without prior knowledge of which Method refers to the deep features or the hand-picked (simple). Using a scale from 1 to 5, 5 being the best, the evaluation metrics used are the following:

1. Global annotations: The annotator defines a score to describe how successful the whole clustering is.
2. Cluster annotations: The annotator defines a score to describe how successful each cluster is.
3. Point annotations: The annotator selects points from different clusters and declares whether they should be approved or rejected in the specific cluster to which they have been assigned. Approximately 100 points were annotated by each user per configuration, for each method.

First we evaluated the global annotations (Figure 3B). For each annotator *i*, we calculated the mean score *μ* of each clustering configuration *s* (KMeans-6, GMM-6, Agg-6), using the scores from the 4 recordings:

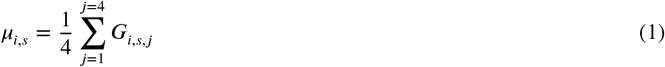

where counter j refers to the 4 recordings and *G_i,s,j_* is the global score of each configuration *s*, in recording *j*, set by annotator *i*. Then, the mean *m* and standard deviation *d* of these mean scores of the 4 users is calculated (Figure 3B).

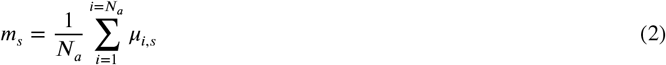

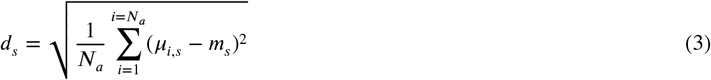

where *N_a_* is the number of annotators (in our case, 4).

**Figure 3.**
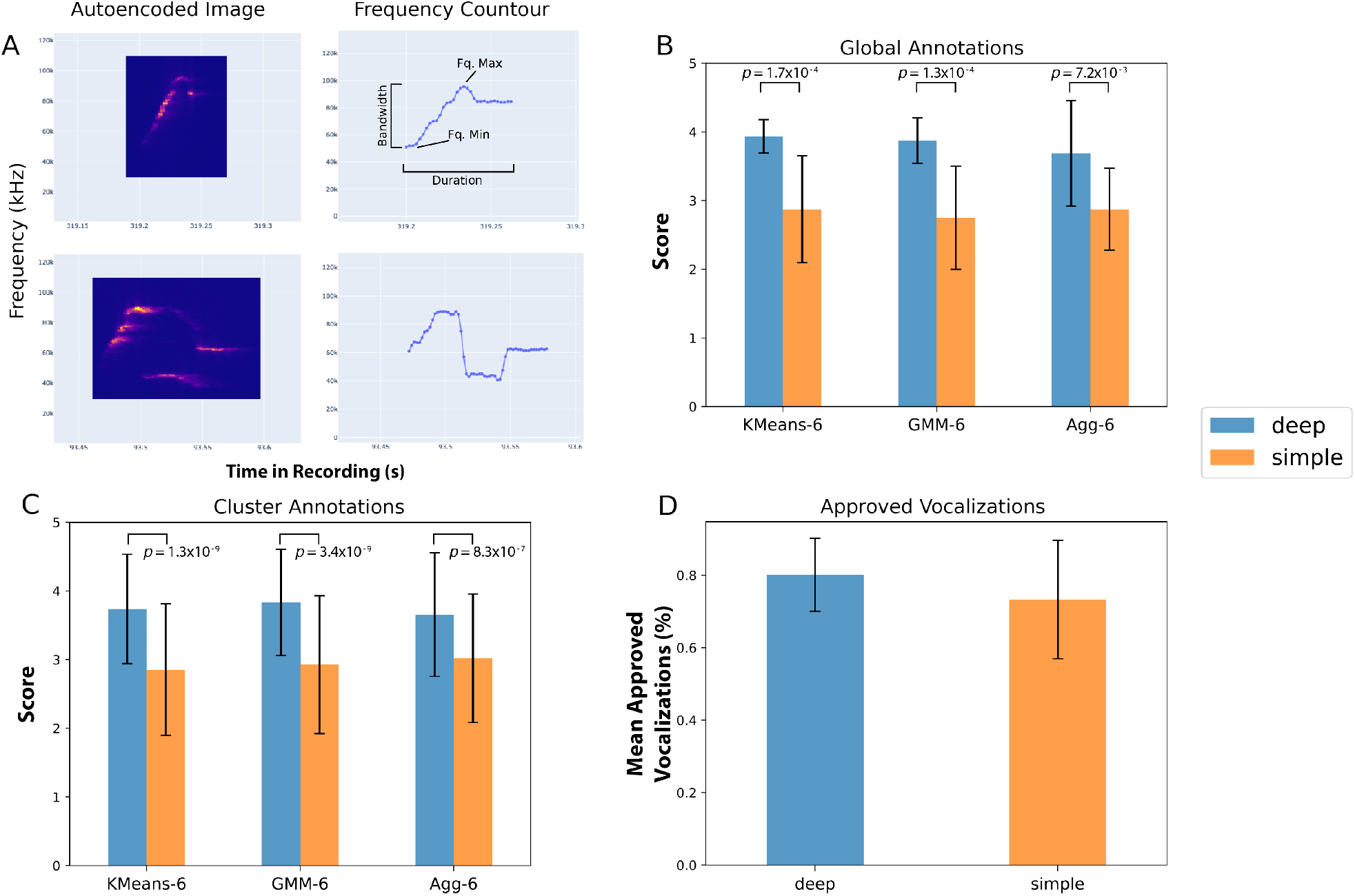
Global and cluster annotation evaluations. A) Examples of two vocalizations with the autoencoder image generated from deep feature extraction (left column) and the peak energy countour used for hand-picked feature extraction (right column). Mean and standard deviation (error bars) of the scores assigned in each configuration for all files evaluated by the annotators for B) the global clustering scores and C) for each cluster in each configuration. D) Mean percentage of approved vocalizations for point annotation evaluation. *p*-values in C and D determined by Student’s *t*-test.

Based on the annotation results, deep feature extraction yields better clustering results than simple feature extraction, with the three configurations used in this test (Figure 3B). All pairs of annotation scores were statistically significant (Student’s *t*-test; K-Means, *t*= 5.0, *p*= 1.7·10^−4^; GMM, *t*= 5.1, *p*= 1.3×10^−4^, Agg, *t*= 3.1, *p*= 7.2×10^−3^). The configuration does not seem to affect the performance of the clustering very much, as far as the mean values are concerned.

Next we looked at the cluster annotations scores (Figure 3C). For each configuration and annotator, we calculated the mean scores *μ*′ of all the 6 clusters in the 4 recordings:

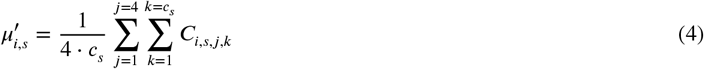

where counter *j* refers to the 4 recordings, *k* to the number of clusters of each configuration *c_s_* (in our case, 6 for all configurations) and *C_i,s,j_* is the cluster specific score of cluster *k* of connfiguration *s*, in recording *j*, set by annotator *i*.

These mean scores were then used to calculate the mean and standard deviation values (*m*′ and *d*′ respectively) for each configuration:

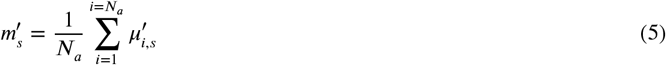

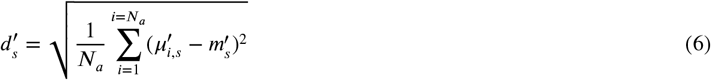

where *N_a_* is the number of annotators (in our case, 4). The results of the cluster-level annotation scores were consistent with the ones from the global annotation scores (Figure 3). As with global scores, all cluster annotation comparisons showed significant differences between deep and simple feature annotations (Student’s *t*-test; K-Means, *t*= 6.7, *p*= 1.3·10^−9^; GMM, *t*= 6.5, *p*= 3.4·10^−9^, Agg, *t*= 5.3, *p*= 8.3·10^−7^).

We then assessed the percentage of vocalizations each user approved for each cluster based on their unsupervised assignment for both deep and simple methods (Figure 3D). If we denote the approved vocalizations of user *i* as *a_i_*, and the total vocalization annotations they have made as *t_i_*, then the mean percentage of approved vocalizations is calculated as:

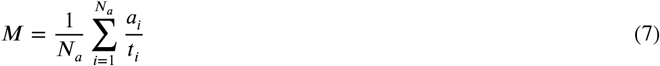

It is clear that users more frequently approved vocalizations clustered by the deep feature extraction method (Figure 3D). Overall, the deep feature extraction method outperforms the simple method in all terms:

- Global clustering evaluation was 37% higher than the simple approach (Figure 3B)
- Cluster-specific evaluation was on average 30% higher than the simple approach(Figure 3C)
- Average point-level evaluation was 10% higher than the simple approach (Figure 3D)

This suggests that the encoder has indeed retrieved useful information of each image, resulting in feature vectors that enable a better clustering of the vocalizations based on visually discernible features.

Further, this indicates that there are much more complex similarities and differences between vocalizations carried by the representations extracted from the encoder than what is available from hand-picked features.

## Discussion

In this paper we have presented Analysis of Mouse Vocal Communication (AMVOC), an open-source tool for detecting and analyzing mice ultrasonic vocalizations. A robust USV detection methodology has been presented and evaluated against a real dataset and results have demonstrated that it is as accurate (and in some cases more accurate) as state-of-the-art tools, while still being fast enough to feasibly function in real-time conditions.

Apart from accurately detecting USVs, AMVOC includes a novel unsupervised approach for representing and grouping the USVs’ content using convolutional autoencoders. We have presented a human annotation procedure for evaluating the ability of the proposed representation technique to discriminate between different types of USVs. Results show that the proposed deep feature extraction method outperforms simple hand-picked features in clustering USVs that are more similar to one another(>30% in global- and cluster-level annotations), leading to possibly meaningful and homogeneous clusters.

This methodology sets a new context for automatic content characterization of mouse vocalizations, according to which clusters of similar vocalizations can be automatically “learnt” through deep neural network architectures. While others have used autoencoders to analyze mouse USVs (***Goffinet et al., 2021***), these methods were not designed to allow users to explore the deep feature clustering and evaluate the results themselves.

In addition to the functionalities and usage modes mentioned, the Dash app in which AMVOC runs provides the user with the ability to save the current clustering configuration, along with other user-friendly functionalities. For example, the user can save the feature vectors and the corresponding clusters of every vocalization, and then use them as ground truth data to train a classifier. By using the automatically extracted vocalization clusters (from the proposed unsupervised pipeline) as classes to train a supervised model, users can then load the model in a real-time and online behavioral context to classify discovered syllables immediately after the vocalizations are detected. This workflow provides a previously inaccessible way to evaluate mouse USV behavior, and opens the door for many real-time, closed-loop behavioral assessments by uniquely utilizing the proposed semisupervised approach. A future direction of the present work is the application of semisupervised methodologies in USVs clustering. The goal would be to integrate human interventions in the classifier used for detection (either offline or online). This could include alternative clusters with reassigned labels for missclustered USVs or comparing pairs of vocalizations, which can be used as pairwise constraints to retrain the encoder part of the model, thus improving subsequent clustering outcomes. These retrained models and new classifications could provide many new opportunities for approaches such as operant behavior, or new types of experiments in mouse vocal research that were previously inaccessible with current methodologies.

While much of the work discussed here and by others focuses on descriptions and characterizations of single USVs, applying these techniques to sequences of USVs could yield a more detailed understanding of rodent vocal behavior. Further development of methods like AMVOC at a sequence-level could provide ways to examine the temporal relationship between syllables across bouts of vocal behavior. It has been known since mouse USVs were first identified as “songs” (***Holy and Guo, 2005***) that across the timescale of a recording session mice will use similar sequences of vocalizations. There have been some attempts to describe the extent to which USV sequencing can be considered patterned (***Chabout et al., 2015***, ***2016***), but these analyses have focused on pairwise changes in sequencing (i.e. how one syllable type follows or precedes another). We propose that future analyses would be best approached at varying timescales and syllable sequence dimensions. By doing so, sequences of vocalizations could be processed and studied, so they can be connected and correlated with various behaviors in mice with a broader appreciation of vocal behavior.

Improvements in, and increased access to high-capacity computing resources that are user friendly and scalable has led to an explosion of resources being developed to measure as many features of animal behavior as is possible (***Datta et al., 2019***; ***von Ziegler et al., 2021***; ***Hausmann et al., 2021***). These resources have become invaluable as neuroscientists push to understand behaviors in more naturalistic contexts. Such unrestrained behaviors can allow for more flexible experimentation and a deeper understanding of ethologically relevant brain function, or clinically relevant behavioral changes (***Jones et al., 2020***). One way that this has been approached in most behavioral experiments is to use markerless animal tracking or pose estimation of single or multi-animal contexts (***Kabra et al., 2013***; ***Mathis et al., 2018***; ***Gal et al., 2020***; ***Marshall et al., 2021***; ***Hsu and Yttri, 2021***). As with mouse USV analyses, many of these methods remain offline, with some notable exceptions providing low-latency feedback for behavioral experimentation (***Kane et al., 2020***). Following in the footsteps of increased data capture and processes that facilitate real-time event detection in animal behavior, AMVOC sets a new standard for vocally-driven analysis of mice behavior.

## Materials and Methods

### Laboratory setup and dataset

We used recordings from two sources: recordings used to develop AMVOC were taken from ***Chabout et al.*** (***2015***), recorded at Duke University. The recordings are publicly available on mouseTube((***Torquet et al., 2016***)). These recordings were captured at 250 kHz. Other recordings used in this study are derived from behavioral experiments detailed below.

#### Animals

Adult C57BL/6J mice were purchased from Jackson Laboratory and a colony was bred and maintained at The Rockefeller University. Before recording experiments, all mice were group housed (2–5 per cage) and kept on a 12-h light/dark cycle, and received ad-libitum food and water. All procedures were approved by the Institutional Animal Care and Use Committee of The Rockefeller University.

#### Behavioral Recordings

Adult male and female mice were used in this study. We used procedures similar to those previously described by Chabout et al. (2015, 2016, 2017). Briefly, before being used for recordings, adult male mice (>8 weeks old) were sexually socialized overnight with a sexually mature female mouse. The next morning, the female mouse was removed and returned to her home cage. For subsequent exposures, when eliciting female-directed ultrasonic vocalizations, different females were used than the one used for sexual socialization. Prior to the vocal recording experiments, the experimental male was removed from their home cages and acclimated in a clean cage inside the recording chamber for 10 minutes. After this acclimation period, either an adult female mouse was introduced into the cage with the male (recordings referred to as FemaleLiveDirected), or urine from a female mouse was placed in the cage (recordings referred to as FemaleUrineDirected). After the female or urine stimulus was introduced, the recording was started. Recording sessions lasted up to 5 minutes. After the recording period was completed, the animals were returned to their home cages. The recording chamber is a 15” × 24” × 12” beach cooler which acts as an acoustically insulated environment. No lighting was provided inside the cooler, as mice are nocturnal and vocalize more in the dark. A microphone was hung from the top of the chamber, placed ~15*cm* from the bottom of the cage. Some recordings were intentionally noisy in order to increase the spectral energy in lower frequencies, thus challenging the detection capacity of AMVOC and other methods. For these “noisy recordings”, we used clean cages containing a bedding of Corn Cob (Lab Supply, Fort Worth, TX) or Cellu-nest™ (Shepherd Specialty Papers, Watertown, TN). For real-time detection, we used an Ultramik 384K BLE (Dodotronics, Italy), sampling at 384kHz, connected to a Raspberry Pi 4 Model B (8GB RAM), and monitored vocalizations via live sonograms using the UltraSoundGate CM16/CMPA recording system (Avisoft Bioacoustics^®^, Berlin, Germany).

#### Datasets

In the course of our experimental and evaluation procedures, we created three datasets (D1, D2 and D3).

Dataset D1 was created to evaluate and compare vocalization detection methods. Specifically, we compiled a ground-truth dataset of 9 audio segments of 5-10 seconds each, containing 245 syllables in total from 14 different mice. The ground-truth annotation was performed by a domain expert by simply declaring the frames that correspond to actual vocalizations, with a time resolution of 1 ms. The recordings used, along with the annotations and instructions on how to reproduce the results, is openly available and can be found in the AMVOC repository https://github.com/tyiannak/amvoc/tree/master.

Dataset D2 consists of 26 different recordings from 9 different mice, used as the training set of our convolutional autoencoder, explained in Section. These recordings can be found under https://drive.google.com/drive/folders/14l-zJmXcjSR9cucnq8lwmUlcsSlRPxe5 and https://drive.google.com/drive/folders/1M976oaxiMpEffN9dfm5Kd5kHeGIiNhbw.

Dataset D3 was created for the experimental evaluation of the clustering configurations (explained in Section)). We used a dataset of 72 behavioral recordings, explained in, 36 in the category FemaleUrineDirected and 36 in the category FemaleLiveDirected. These recordings come from 12 different mice. We have randomly selected 20 s from each recording, where the vocalization rate should be at least 2.5 vocalizations/sec. We then concatenated the 20 seconds interval from each recording to a new recording. We generated 4 recordings, 2 from the FemaleUrineDirected category and 2 from the FemaleLiveDirected category. These recordings can be found under https://drive.google.com/drive/folders/1l7qUw0SVvd1dzNr35FT7XOxbhxDp7Kqn.

### Detection of mouse vocalizations

The first step of AMVOC’s processing pipeline is to detect mice USVs. Given an input recording, we want to determine the parts of the signal that contain a USV. Below are methods for offline and online detection.

#### Offline USV Detection

In order to detect the mice USVs, we first compute the spectrogram of the whole recording. This is done by splitting the signal to non-overlapping short-term windows (frames) of duration *w* = 2 ms (time resolution) and calculating the Short-Term Fourier Transform (STFT) for each time frame. Frequency resolution *f_r_* is calculated as:

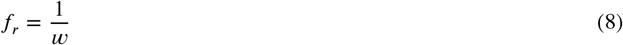

This means that in our case, if *w* = 2 ms, *f_r_* = 0.5 kHz. In simple terms, the ability of the 2-dimensional (time & frequency) spectrogram representation to discriminate between different frequency coefficients is 0.5 KHz (i.e. frequency resolution). This is less fine-scaled than is typical for human speech analytics methods, but because mice vocalizations usually range in the frequencies 30-110 KHz, this resolution is more than sufficient.

As soon as the spectrogram is extracted, the USVs are detected on a time frame-basis, using two separate criteria, time-based thresholding (TT) and frequency-based thresholding (FT), that take into account the values of the distribution of the signal’s energy at the different frequencies (Figure 4A). Both of these criteria are based on the spectral energies, however they differ in the way the thresholding criteria are calculated and applied. The details of the two criteria are as follows:

- Time-based thresholding (TT): This involves a simple temporal thresholding of the spectral energy values. To do this, for each time frame, we calculate the spectral energy by summing the energy value at each frequency. We do this procedure for the frequency range of interest, which is, as mentioned above, from 30 kHz to 110 kHz. If we denote the spectrogram value at time frame *i* and frequency *j* as *E_ij_*, spectral energy *S_i_* is calculated as:

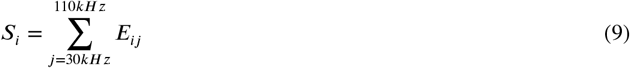

where the step of *j* is equal to 0.5 kHz. We then compute a dynamic sequence of thresholds for the spectral energy. In particular, for each frame *i*, for which we have extracted the spectral energy *S_i_*, we compute the dynamic threshold:

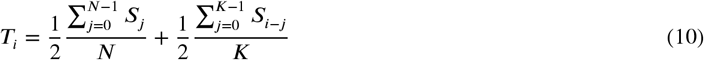

where *N* is the number of time frames, so the first term refers to the mean spectral energy. In the second term, *K* is the size of a moving average filter in seconds. Here we use *K* = 2 seconds, which is convolved with the sequence of spectral energy values. In other words, the dynamic threshold *T_i_* is defined at each frame *i* as the average of the current spectral energy (*S_i_*) and the moving average of the spectral energies of the last *K* frames.
- Frequency-based thresholding (FT): This second criterion is associated to applying a thresholding rule, based on the per-frame distribution of energies on the different frequencies. A simple dynamic threshold at each time frame of the spectrogram (as described above) is not enough, because there are also time frames where high spectral energy occurs due to noise. The spectral energy value in these high-noise time frames may surpass the threshold, but this does not correspond to any vocalization (Figure 4B and C). Our goal is to filter out these false positive vocalizations. It is easy to observe that in the vast majority of cases, noise appears as high energy values, spread across a large frequency range in each time frame (Figure 4C), compared to the time frame energy distribution in vocalizations, which is concentrated in a small frequency range in each time frame (Figure 4B). Our filtering criterion was to keep only time frames where the peak energy value *P_i_* is larger than the mean spectral energy (*M_i_*) of a 60kHz range around the frequency of the peak energy (truncated if the range goes below 30kHz or above 110kHz). If we denote the energy value at time frame *i* and frequency *j* as *E_ij_*, the equations describing the two quantities above, are the following:

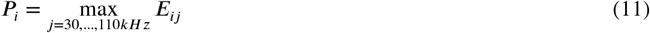

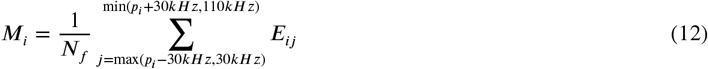

where the step of *j* is equal to 0.5 kHz and, as a result, *N_f_* = 2 · (min(*p_i_* + 30*kHz*, 110*kHz*) − max(*p_i_* − 30*kHz*, 30*kHz*)), and *p_i_* is the frequency of the peak energy at time frame *i*, i.e. 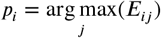.

**Figure 4.**
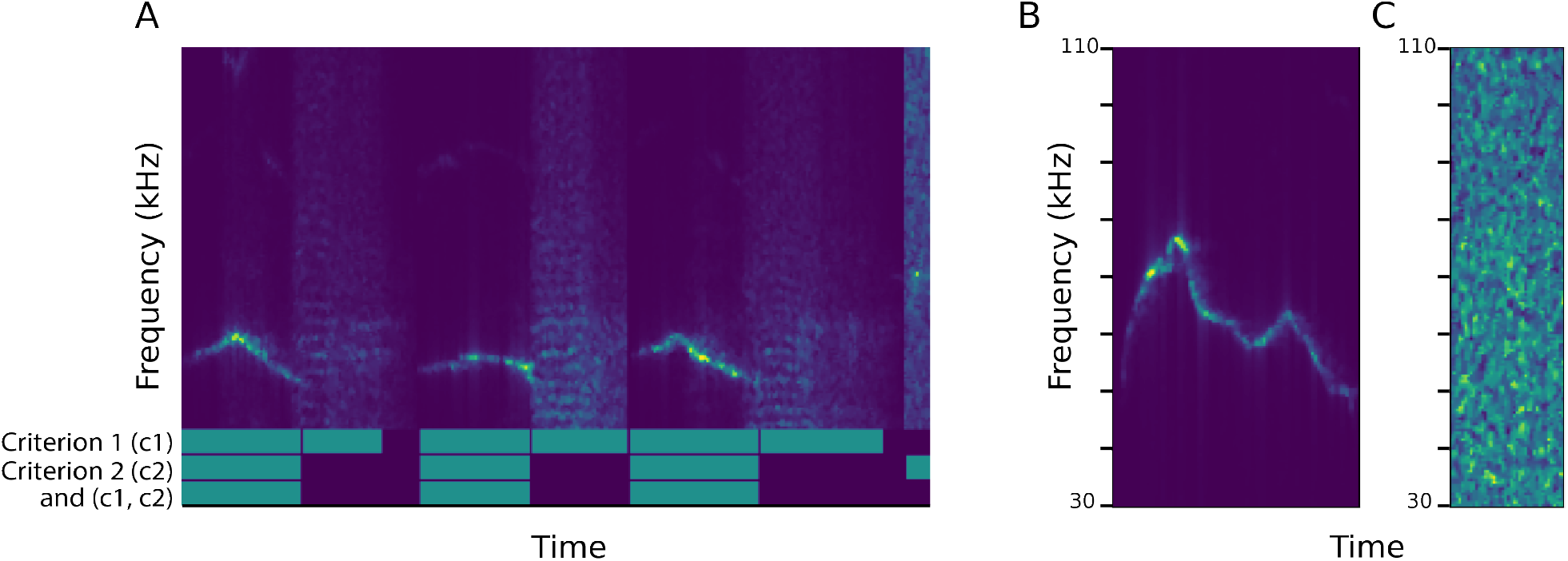
Examples of detection criteria. A) Demonstration of the twofold thresholding application. The green bars of the first two lines show the detected vocalizations by each criterion, whereas the third-line green bars are their intersection. Segments were spliced for purposes of visualization. B and C) Examples of two different segments of the spectrogram, from 30-110 kHz. Here, an actual vocalization (B) and a noisy, hihg-energy segment are displayed. In the noisy segment each time frame surpasses the spectral energy threshold (criterion 1), but not the second applied threshold (criterion 2).

Both criteria TT and FT are applied on each short-term frame *i* as follows: the threshold conditions require that the spectral energy is higher than 50 percent of the dynamic threshold computed in step 1 and that the maximum energy is larger than the mean spectral energy by a factor of 3.5. Let *V* be a sequence of frame-level vocalization decisions, i.e. *V_i_* = 1 if time frame *i* is part of a vocalization and *V_i_* = 0 if not (Figure 5A). Then the above rule can be expressed as follows:

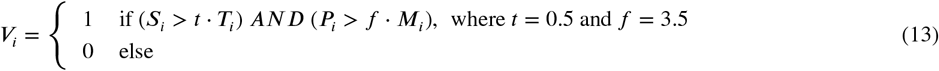

**Figure 5.**
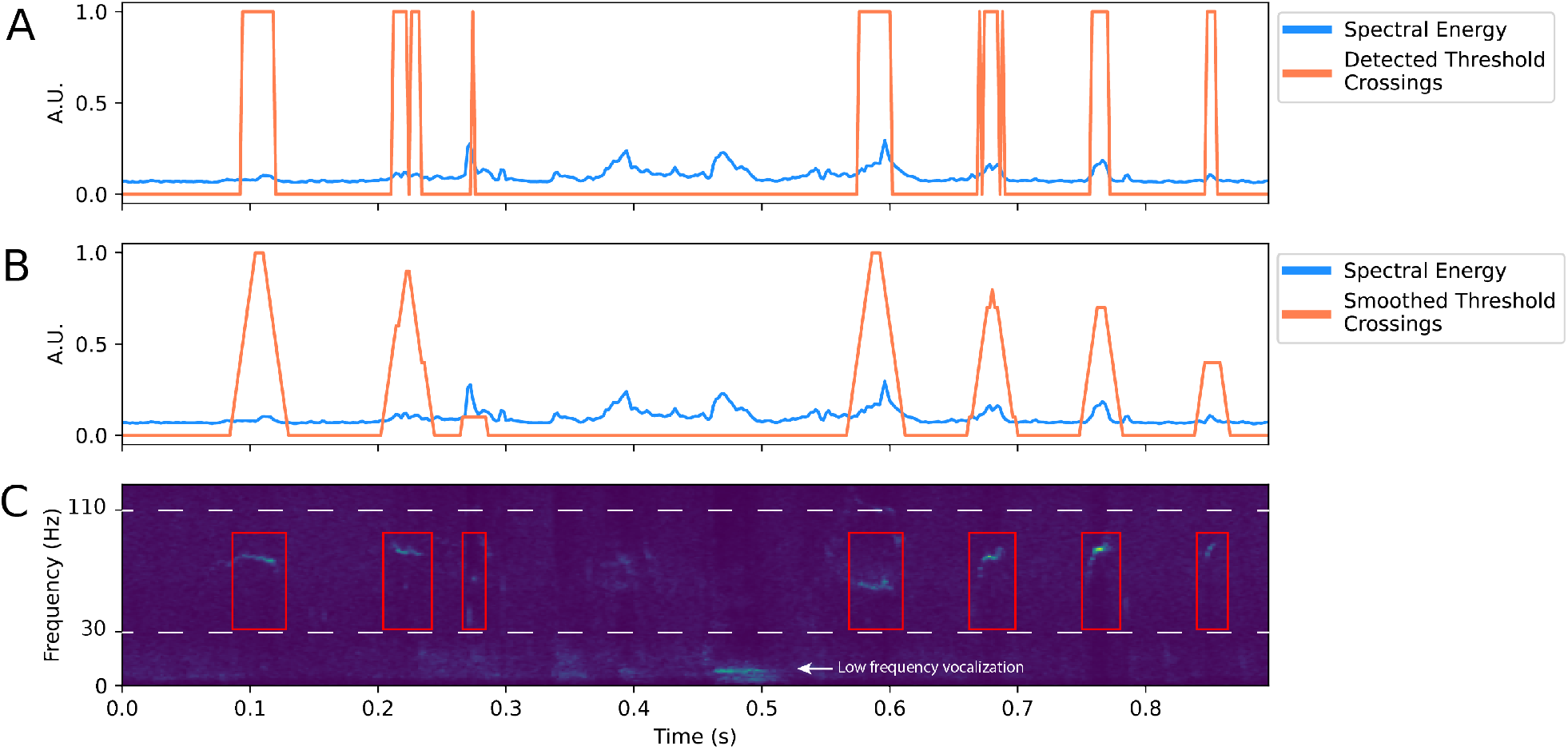
Detection by spectral energy threshold and smoothing. A) When spectral energy (blue) crosses a particular threshold the start and end of a vocalization is determined (orange). B) Threshold crossing is smoothed to determine final event and temporal detection (bottom). C) Representative spectrogram indicating where the USVs were detected based on threshold crossings. Dotted white line denotes 30-110kHz boundary used for detection.

Both factors *t* and *f* have been selected after experimentation and can be considered as configurable (see Section).

After the twofold thresholding rule has been applied as described above, we apply a smoothing step. Specifically, after the sequence *V* of 1s and 0s occurs (Figure 5A), this sequence is smoothed using a moving average filter with a duration of 20 ms (Figure 5B), so that the neighborhood of the possible vocalization is taken into account:

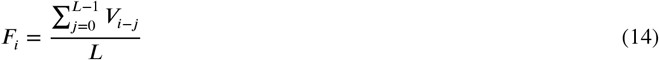

where is the size of the filter. As a final step, if successive positive frames found in *F* are separated by <11 ms, they are concatenated to form segments of mice vocalizations (Figure 5C). Vocalizations of duration <5 msec, are filtered out, since practical evaluation showed that in most cases these are false positive detections.

#### Online USV Detection

In addition to the offline detection that is standard for most USV analyses, we also developed an online version of AMVOC (i.e. using streaming sound recorded from the computer’s soundcard). This is achieved by following the aforementioned analysis steps, though these cannot be applied to the whole signal at once, since, in a real-time setup, the signal is recorded simultaneously with the detection procedure. For this process, we wanted the processing interval to be as small as possible, in order to detect the vocalizations fast. On the other hand, if we process the signal more often, the probability of cutting a vocalization in the border between two successive blocks is increased. Additionally, the signal statistics that need to be calculated for the detection steps described in the preceding section will become less robust if they are computed on smaller segments. We therefore chose to process the signal in blocks of a fixed duration, which for our case has been set equal to 750 ms. We estimated the number of USVs that could occur per processing window by running 20 recordings from live female and 20 recordings from female urine social contexts by processing the files with the online computations of AMVOC in an offline format. A 750ms window provides a long enough period for high accuracy detection and minimally interrupting the number of individual syllables that would be processed (Figure S1A and B). AMVOC requires about 5ms to process a 750 ms window (Figure S1C). This processing time is independent of the number of USVs per window, and appears constant across tested files (Figure S1D and E). Since this processing time is less time than any inter-syllable interval (Figure S1F), this means that no USVs are lost to drop-out due to computational load or processing time.

The main algorithmic difference between the online and the offline detection is the calculation of the dynamic threshold. If we denote *k* as the current block, the dynamic threshold is the same for all frames belonging to that block and it is computed as follows:

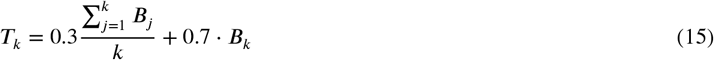

where *B_k_* is the mean spectral energy of block *k* in the 30-110 KHz frequency range (as described in Section). In simple terms, the block’s threshold is computed as the weighted average of the mean of the spectral energies of the blocks recorded up to that point and the current block’s spectral energy. The weights (0.3, 0.7) were chosen after extensive experimentation (Figure S2). The sequence of thresholds that occurs is then multiplied by the threshold percentage *t*, exactly as in the offline method (Section). Another difference between the offline and the online methodologies is that for the online approach, we have selected to use an overlapping block. In particular, we always process the newest 750 ms recorded segment plus the last 100 ms of the previous block. In this way we add a minor computational delay (as we repeat the process for 13% of the data), and we manage to eliminate the errors caused by USVs being split between two successive blocks, and therefore lost by the detection method. The rest of the procedure is the same as the offline method and it is applied every 750 ms.

#### Vocalization Detection Configuration

As mentioned earlier, the two parameters which determine the vocalization detection procedure is the threshold percentage *t* and a factor *f*; this would mean that energy of a 60 kHz area *M_i_* around the frequency of peak energy *p_i_* must surpass peak energy *P_i_* by the factor *f*. The user can change these values according to the expected recording conditions and application requirements. Parameters used in our current study were optimized to include small events while minimizing false positives. To select these parameters, we used Dataset D1 (Figure 6). As expected, increasing either of these parameters results in a more strict thresholding, which means that the precision increases, and recall decreases. The opposite is observed when either of the parameters is reduced. From a more qualitative point of view, increasing the threshold might result in splitting a vocalization with relatively low peak energy in intermediate time frames. On the other hand, a very low threshold can merge successive vocalizations.

**Figure 6.**
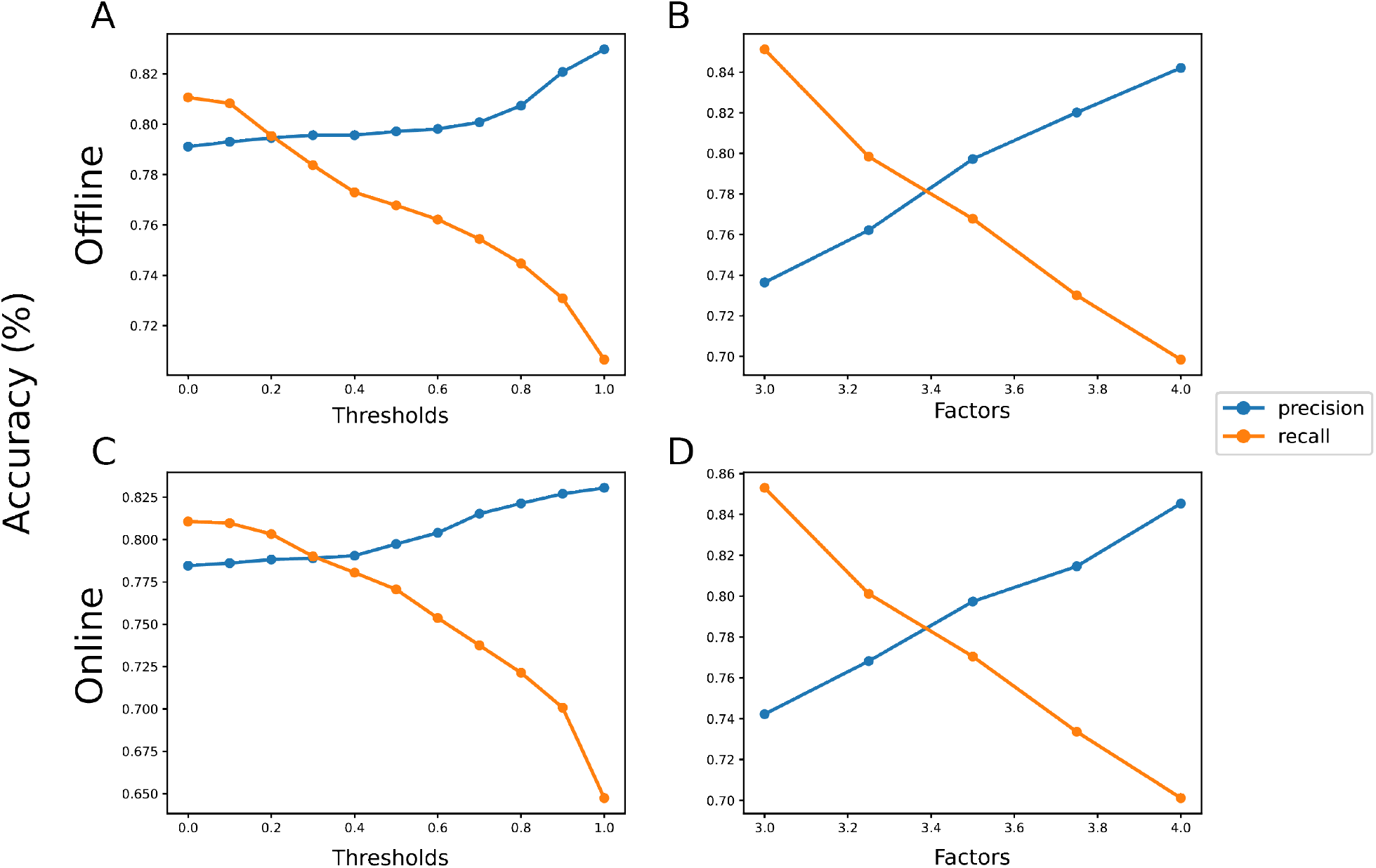
Effect of changing parameters on precision and recall. A and B) Changes in precision and recall during offline detection when thresholds *t* and factors *f* are changed. C and D) Changes in precision and recall during online detection when thresholds and factors are changed. Optimal threshold and factor were determined to be the same, 0.5 and 3.5, respectively, in both detection modes.

### Deep unsupervised learning for mouse vocalization clustering

We next developed a method for unsupervised clustering of the detected vocalizations, by using a convolutional autoencoder, from which features are derived.

#### Convolutional autoencoders

##### Autoencoders

Autoencoders are neural networks that are trained to attempt to copy their input to their output. An autoencoder consists of 3 components: encoder, code and decoder. The encoder compresses the input and produces the code, the decoder then reconstructs the input only using this code. The encoder is described by a function *h* = *f* (*x*), where *x* is the input and *h* is the code and the decoder produces a reconstruction *r* = *g*(*h*).

The encoder maps the input into a code that is assumed to contain the most important information (features) of the input, and then decoder constructs the output from this representation. The intermediate representation of the input, is also called the latent-space representation.

By training the autoencoder to perform the input copying task we hope that the code *h* will be an efficient representation of the input and will contain important information from it. One way to obtain useful features from the autoencoder is to constrain *h* to have a smaller dimension than *x*. An autoencoder whose code dimension is less than the input dimension is called undercomplete. Learning an undercomplete representation forces the autoencoder to capture the most salient features of the training data. The learning process is described simply as minimizing a loss function *L* (*x, g*(*f* (*x*))), where *L* is a loss function penalizing *g*(*f* (*x*)) for being dissimilar from *x*.

Commonly, the encoder and the decoder are feedforward neural networks, whose task is to learn the function *f* and *g* respectively.

In our case, where the inputs are images, the encoder and the decoder are convolutional neural networks.

#### Convolutional neural networks

CNNs are a kind of network that process data with a known grid-like topology, like image data. They are named after the mathematical operation convolution, which they employ. Their fundamental characteristics, which we make use of, are:

- 2D convolution: The goal of a 2D convolution is to produce the activation map. More specifically, we have the input image and a specific number of filters, which are kernels of fixed dimensions, e.g 3×3. Every filter corresponds to a unit of the network, which is connected to a certain e.g. 3×3 area of the image (receptive field) and learns its features and properties. So, the 2D convolution is responsible for learning the local characteristics of the image. Each kernel is convolved with the whole image and produces a convolutional activation map. After the convolution, we will have activation maps of depth equal to the number of the filters applied.
- Pooling: It is performed after each convolution and reduces the size of the activation maps. This procedure is important, because the computational load of the next layers of the network is reduced, since the width and height of the convolutional activation maps is downsampled. This enables us to obtain a representation of the image which is scale-independent. By reducing the size of the image after every convolution, we ensure that the intermediate representation will indeed have lower dimensions compared to the initial image.

#### Applications

Autoencoders are powerful tools for dimensionality reduction or feature learning. Applications of undercomplete autoencoders include compression, recommendation systems as well as outlier detection. Convolutional autoencoders are frequently used in image compression and denoising. They may also be used in image search applications, since the hidden representation often carries semantic meaning. Recently, theoretical connections between autoencoders and latent variable models have brought autoencoders to the forefront of generative modeling. More specifically, the variational autoencoder is a generative model which can be trained and used to generate images (***Goodfellow et al., 2016***).

In our case, the purpose of adopting a convolutional autoencoder is to create meaningful feature exctractors for the spectrograms that correspond to the detected mice vocalizations. The input of the autoencoder are the images (spectrogram parts) that contain a vocalization and our goal is to obtain an intermediate representation (code *h*), in order to use it as a feature vector that will uniquely describe each image. From this representation, the image will be reconstructed in the decoder and the output will be compared to the input in each step, in order to calculate the loss function. The structure and the training of our autoencoder is described below.

#### Proposed autoencoder architecture and training

To train the autoencoder, we used Dataset D2 (see Section) and we calculated the spectrogram for each of its recordings. This dataset contains 22,409 detected syllables. Each vocalization is represented by a spectrogram in the detected time interval and defined frequency range. Therefore, these spectrograms vary in width, which corresponds to the respective syllable duration. So, we have to specify the width of the images that we are going to feed to the autoencoder, since the frequency y axis is the same for all spectrograms: 80 kHz range (from 30-110 kHz) and 0.5 kHz frequency resolution, resulting in a dimension of 160. Selecting a fix sized time dimension for our spectrograms requires taking into consideration a tradeoff between losing important information from the larger spectrograms (if we crop them) and reducing the importance of the shape and details of the smaller spectrograms (if we zero-pad them), which are more numerous than large spectrograms (Figure 7).

**Figure 7.**
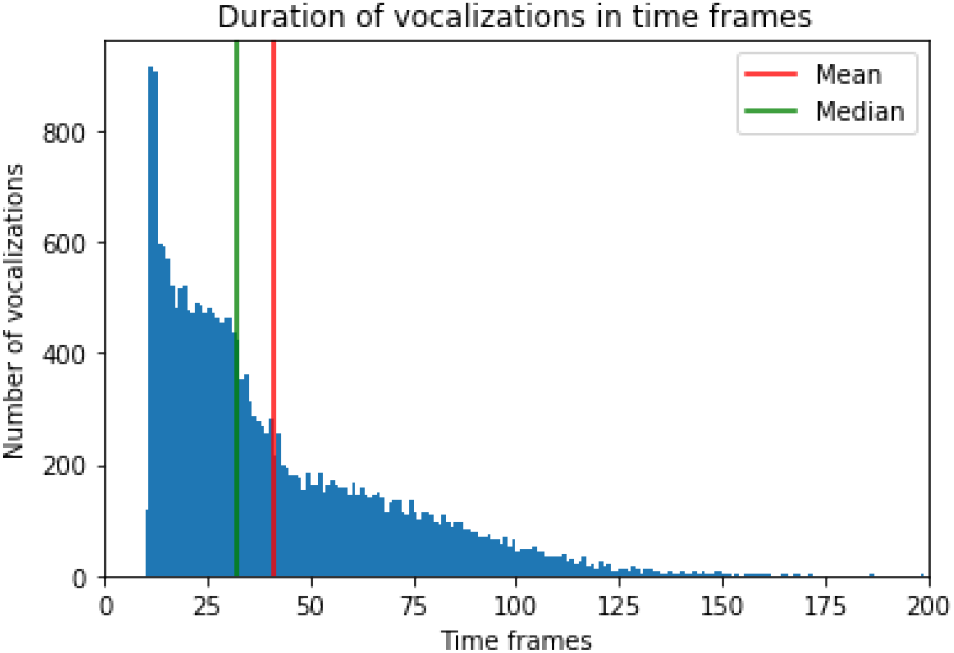
Histogram of the duration of the vocalizations in time frames Each time frame corresponds to a 2 ms duration.

In order to decide the final fixed size of the spectrograms to feed the autoencoder, we plotted a histogram of the initial durations of all the detected syllables in the training set (Figure 7). To ensure uniform sizing, we zero padded small spectrograms, and cropped larger spectrograms keeping the central part of the image. Based on the histogram we selected a fix-sized duration of 64 windows since it is larger than both the mean and the median of the durations (Figure 7). Using this size, we noted that it balanced the information tradeoff mentioned above. It is also a power of 2, which is convenient for the pooling operations in the autoencoder. This length of 64 frames corresponds to 64 time frames × 0.002 sec/time frame = 0.128 sec = 128 ms. The aforementioned process of cropping or expanding spectrograms to a fix-sized width of 64 windows leads to spectrograms of a final resolution of 64 time frames × 160 frequency bins.

The training set consists of 22,409 images, each one with size 64×160. We fed the images to the encoder, which is a convolutional neural network with 3 convolutional layers, each followed by a max pooling layer (Figure 8A). The first convolutional layer uses 64 filters, with dimensions 3×3 each.

**Figure 8.**
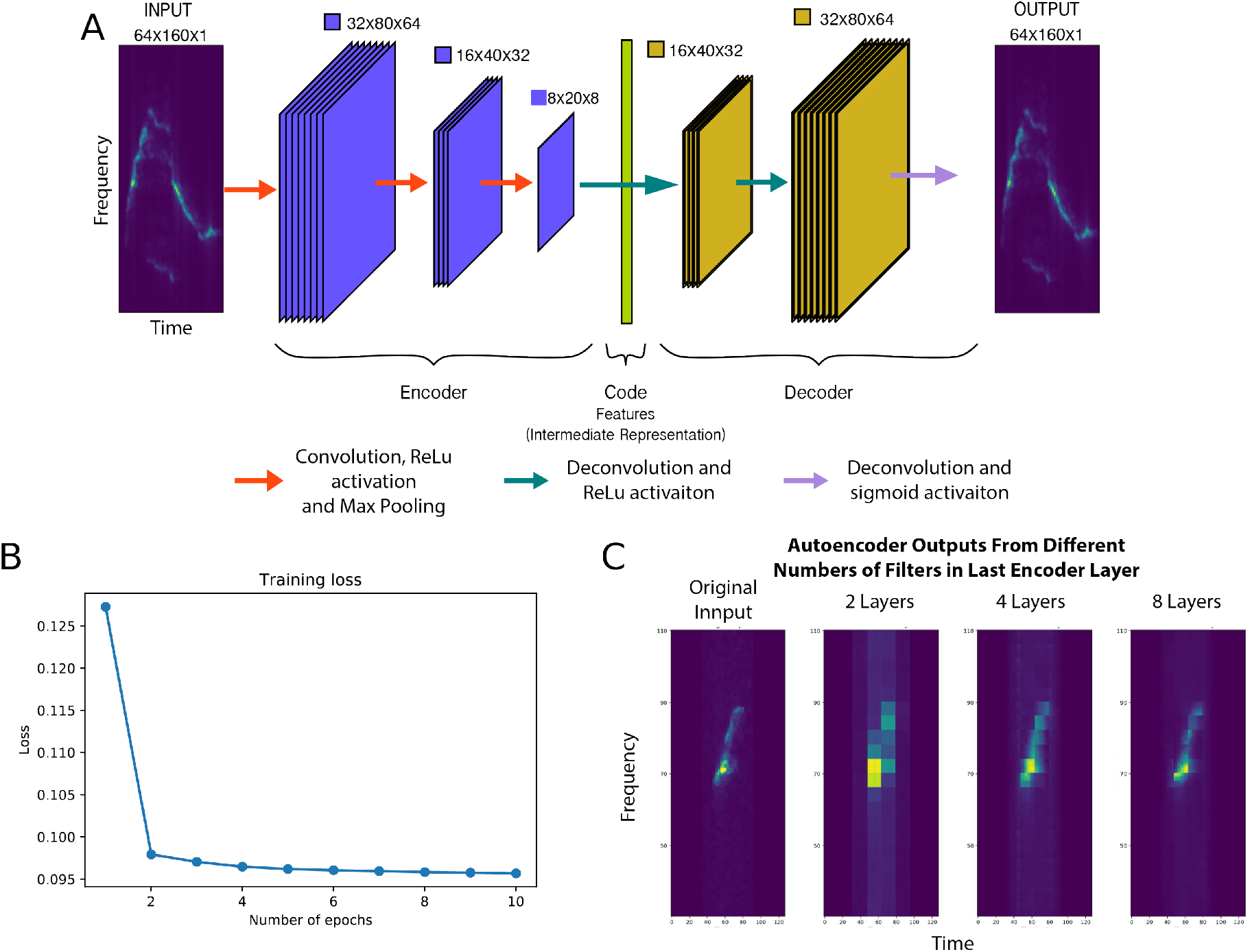
AMVOC convolutional autoencoder A) Architecture used for the autoencoder in AMVOC. B) Effect of the number of training epochs on measured training loss. C) Examples of image reconstruction with AMVOC’s autoencoder after training, using 2, 4 and 8 filters in the encoder output layer. Data is extracted from the input image (left) and used to reconstruct the three images (right).

After that, a max pooling layer decreases the spatial dimensions of the images by a factor of 2. This means that the output of the max pooling layer is a 32×80×64 representation. The next convolutional layer consists of 32 filters, with a max pooling layer generating an output with dimensions 16×40×32. The third and final layer includes 8 filters, and a max pooling layer, resulting in a convolutional activation map for each image with dimensions 8×20×8. This flattened intermediate representation is the feature vector that uniquely describes the input image (i.e. the code).

In order for the convolutional encoder to be trained, and because the task is unsupervised, the second part of the autoencoder (the decoder) is responsible for reconstructing the image we fed to the encoder from the intermediate representation (Figure 8A). The decoder reverts the steps of the encoder, using 32, 64 and 1 filter in the last layer. In each decoder layer, we use filters of size 2×2 and a stride of 2, so that after each layer the size of the representation increases by 2 and the final output of the autoencoder is an image with the same size as the original input.

After each convolutional layer, the ReLU activation function is used, since we want the activation maps to consist of positive values. An exception is the last deconvolution of the decoder, where the Sigmoid activation function is necessary for the reconstruction of the image and the calculation of the Binary Cross Entropy Loss function, as it outputs a value between zero and one.

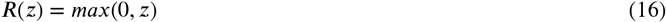

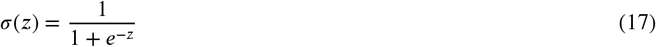

The loss function is calculated based on the differences between the reconstructed and the original image. This loss function indicates the quality of the autoencoder’s performance, and can be improved with increased training epochs. If we denote the output of the decoder at a specific pixel as 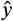 and the actual value of the input image as *y*, then Binary Cross Entropy Loss is calculated as:

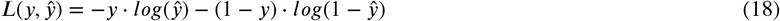

#### Parameter tuning

As described above, the basic parameters for configuring our proposed autoencoder procedure are the following:

- *Number of layers of the encoder*. We tested a range of different numbers of layers (2-4 layers). Using 2 layers appeared problematic, since the autoencoder couldn’t clearly reconstruct input images. Four layers or more result in too many parameters, which slowed down the training, resulting in a bigger loss, while the final reconstruction wasn’t better than the one obtained from a 3-layer autoencoder (Figure S3A).
- *Number of filters per layer* We used the most filters in the first layers (as is typical for classical neural networks), and reduced the number of filters as we went deeper in the network. We tested the autoencoder with varying numbers of layers. Fewer filters resulted in losing information from the images, while more resulted in too many parameters to be trained. The critical choice we had to make was the number of filters in the last encoder layer, the output of which we use as the representation for the specific image in the clustering task. We tested 2, 4 and 8 filters. Using 8 filters resulted in smaller loss, as expected, although using 4 filters were enough for the reconstruction of the images (Figure 8C), meaning enough features for the reconstruction were extracted with fewer filters. We selected 8 filters in our final design in order to ensure that all the details of the various shapes of the vocalizations are properly extracted.
- *Filter size* We experimented with 3×3, 5×5 and 3×5 kernels. A 3×5 kernel appeared reasonable because of the non-square shape of the images, but gave similar results to 3×3 kernels, so we selected the latter, since symmetric kernels are more commonly used (Figure S3B).
- *Size of max pooling kernels* We experimented with reducing the image size by 2 in both dimensions, or by 2 in time dimension and by 4 in frequency dimension due to the non-square image. A 2x symmetrical reduction provided better results (Figure S3C).

As far as other hyperparameters are concerned, we used the Adam optimizer, with learning rate equal to 0.001 and batch size equal to 32.

Training epochs were determined experimentally. We found that 2 or 3 epochs was enough for a good reconstruction of the images, as loss did not decrease much after 3 epochs (Figure 8B). We also did not want to overfit to the training data. Thus, we elected to train the model for just 2 epochs.

An example of the input and output of the autoencoder is shown in Figure 8C. The input comes from a recording that was not used in the training Dataset D2. The reconstruction is lossy, due to our use of an undercomplete autoencoder.

#### Feature extraction and clustering

After the model has been trained in the unsupervised manner described above, it is ready to be used in the feature extraction procedure (Figure 2A). An audio file is selected and its spectrogram is calculated, and individual USVs are detected. The raw spectrograms of the USVs are fed to the autoencoder in batches of 32 and the intermediate representations are derived. These are the feature vectors. Each flattened feature vector has a dimension of 1,280 (8×20×8) after a dimensionality reduction from a dimension of 10,240 in the initial flattened vector, since each image started with a shape of 64×160.

After the features are extracted, we further reduced the dimensions by excluding features that have the smallest variance and will likely be less impactful for the discrimination of USVs. After extensive qualitative experimentation, assuming we have *N*-dimensional feature vectors, we selected a threshold equal to:

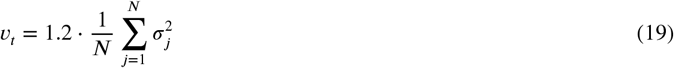

where 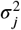 is the variance of feature *j*. In this case, *υ_t_* is the mean of the features’ variances; features with variance less than *υ_t_* will be excluded, resulting in feature vectors undergoing a dimensionality reduction of a factor approximately equal to 4.

Next, we normalize the features using a Standard Scaler. Each feature is scaled according to the following equation:

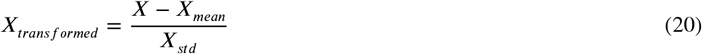

where *X_mean_* is the mean of the feature values for all samples and *X_std_* is the standard deviation of the feature values for all samples.

In the last preprocessing step, we used PCA to further reduce the dimensionality of the feature vectors. Since we do not know the number of components beforehand, we choose the smallest number of components which maintains 95% of the variance of the features before the PCA. Overall, our goal was to both extract many features from the images using the encoder, so that details of the images are taken into account, and simultaneously reduce them as much as possible by ignoring the non-significant features. This final reduced feature representation is then used for clustering. The final visualized cluster is plotted as a 2-dimensional tSNE reduction of the data (Figure 2B).

Each cluster should consist of vocalizations that share some common features that allow them to belong in the same group. Since we wanted the user to be able to choose the number of clusters, we chose clustering algorithms that can have this parameter predefined. Users can choose between the following clustering methods: Agglomerative, Birch, Gaussian Mixture Models, K-Means and Mini-Batch K-Means. In the implemented GUI (see Section), the user can choose one of these clustering methods and the number of clusters, in a range from 2 to 10.

### Baseline feature extraction

To evaluate the quality of AMVOC’s deep feature extraction and clustering, we compared clustering on deep features against clusters derived from hand-picked acoustic parameters (Figure 3A). The hand-picked features were measured as follows:

1. We first calculate the spectrogram in the specific time segment that corresponds to the vocalization and in the defined frequency range (30-110 kHz).
2. We then perform frequency contour detection:
  - Detect the position and the value of the peak energy in each time frame. If we denote the spectrogram value at time frame *i* and frequency *j* as *E_ij_*, the equations describing the two quantities above, are the following:

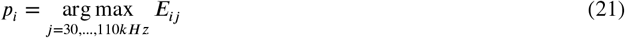

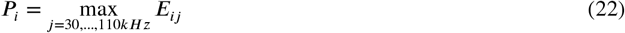
  - Use a thresholding condition to keep only the points *i* where the peak energy is higher than 20 % of the highest energy value in the specific time interval:

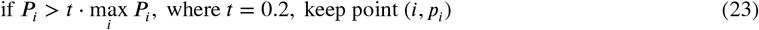
  - Train a regression SVM to map time coordinates to frequency values, using the chosen points (*i, p_i_*) as training data.
  - Predict the frequencies for the same time range. After that, for each vocalization, a frequency contour *c* is created, along with a corresponding time vector *v*, which matches every frame *i* to its actual time of occurrence. This estimated sequence *c* captures the “most dominant” frequency in each time frame, so we can think of it as a spectral shape sequence of each mice vocalization.
3. After the frequency contour *c* is produced, we proceed to the feature extraction step. We selected 4 different features, all based on the frequency contour:
  - Duration of the vocalization *d*. If we denote the number of frames of which the vocalization consists as *N*:

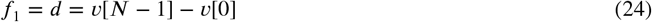
  - Time position of the minimum frequency (of the predicted frequencies), normalized by the duration of the vocalization:

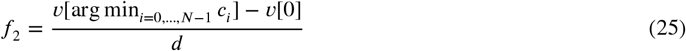
  - Time position of the maximum frequency (of the predicted frequencies), normalized by the duration of the vocalization:

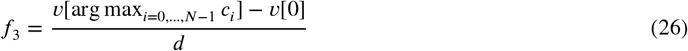
  - Bandwidth is calculated as the difference between the first and the last predicted frequency value, normalized by the mean frequency of the vocalization *m_f_*:

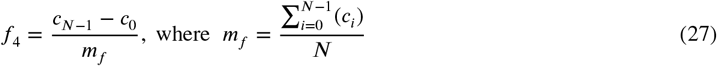

If we interpret the frequency contour as a 2-dimensional graph, where the x-axis corresponds to time and the y-axis to frequency, the 2nd feature is the x-position of the minimum of the curve, the 3rd feature is the x-position of the maximum of the curve and the 4th feature is the normalized difference between y-positions of first and last point. For example, the last feature can discriminate contours with different slopes. After the features are extracted, we scale them using a Standard Scaler (see Equation 20).

### Implementation and library description

The functionalities described throughout the paper were implemented in Python 3.8. The code can be found in the repository https://github.com/tyiannak/amvoc/tree/master/data/vocalizations_evaluation. For the implementation of the autoencoder we used the Pytorch deep learning framework, and we used the clustering algorithms implementations from Scikit-Learn. The visualization of the data in the offline mode, including the clustering, specific vocalizations, and the evaluation choices are all displayed and accessible in the Dash GUI.

## Acknowledgements

We thank E. Waidmann, R. Agravat, and other members of the Jarvis lab for their thoughtful conversations on this project. We thank J. Chabout for providing open access to his past mouse recordings.

## Notes on contributor(s)

VS, CDMV, PFS, and TG co-wrote the paper, and EDJ edited the paper. VS and TG wrote the code for AMVOC. PFS developed MSA2. VS, CDMV, and TG contributed to analyses. CDMV performed vocal behavior experiments and tests. TG and EDJ co-supervised the study.

## Supplementary Material

**Figure S1.**
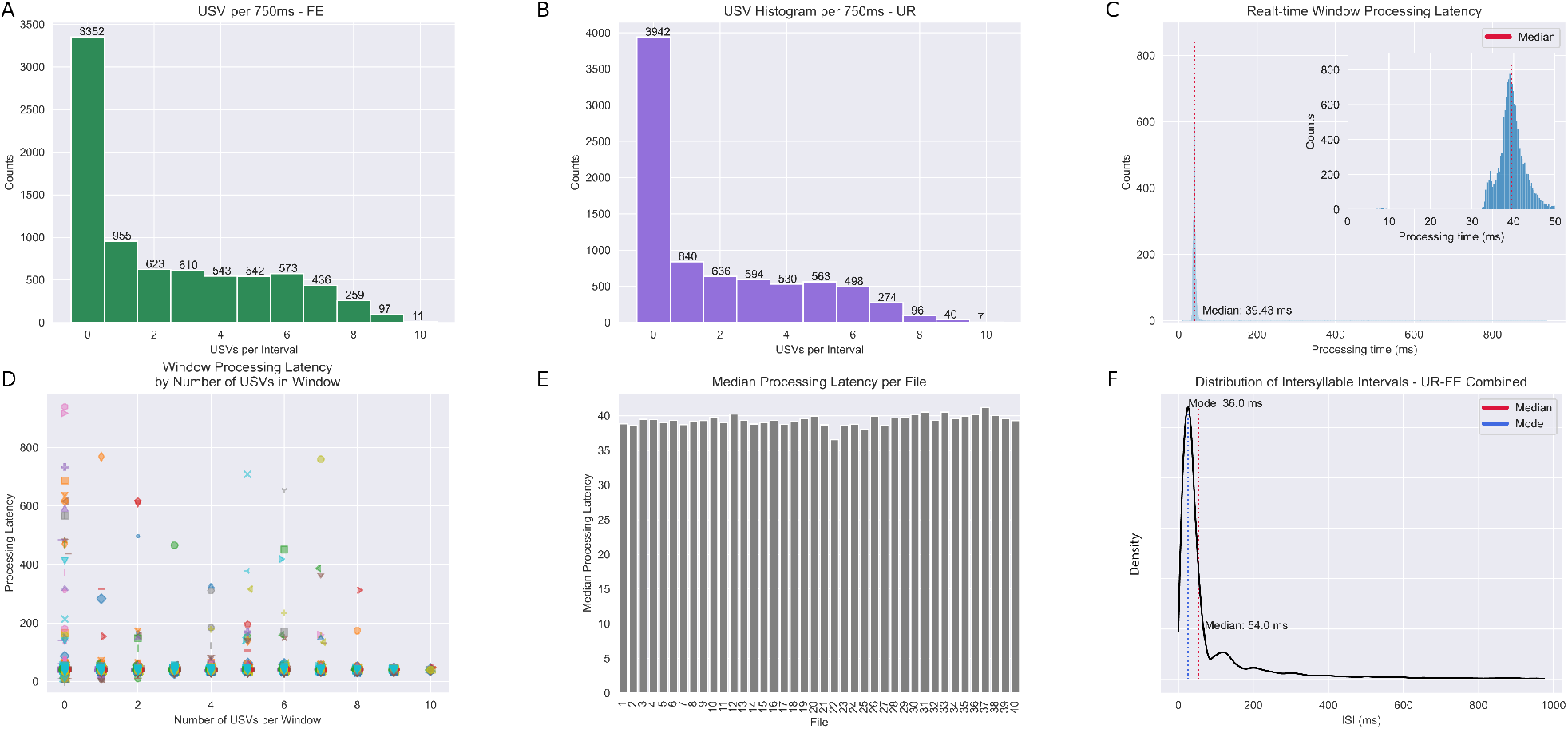
Frequency histograms from real-time processing windows. A) Number of USVs per 750ms non-overlapping segment from 20 recordings of male mice in a live female context. B) Number of USVs per 750ms non-overlapping segment from 20 recordings of male mice in a female urine context. C) Processing time required for each 750ms segment, mean and median are less than 250μs. Inset represents bins from 0-10ms of processing latency. Dashed line indicates median processing time. D) processing latency per number of USVs in a 750ms window. Symbols represent windows within a file. E) Median processing latency of all windows in tested files. F) Distribution of inter syllable intervals among USVs detected in test files.

**Figure S2.**
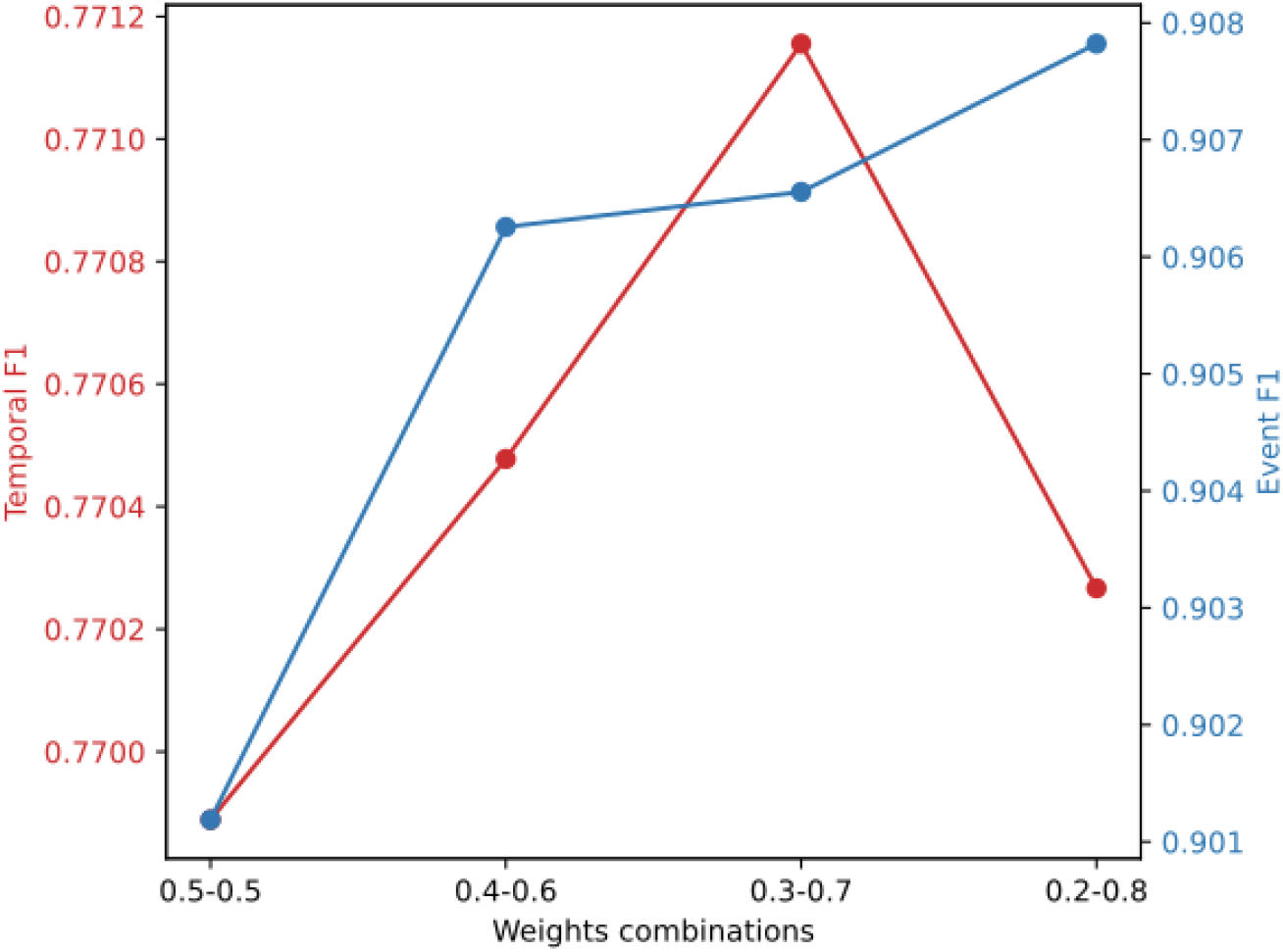
Effect of different weight combinations on Temporal and Event F1 Scores) Temporal (red) and event (blue) accuracy of USVs from our ground truth dataset using different combinations of weights.

**Figure S3.**
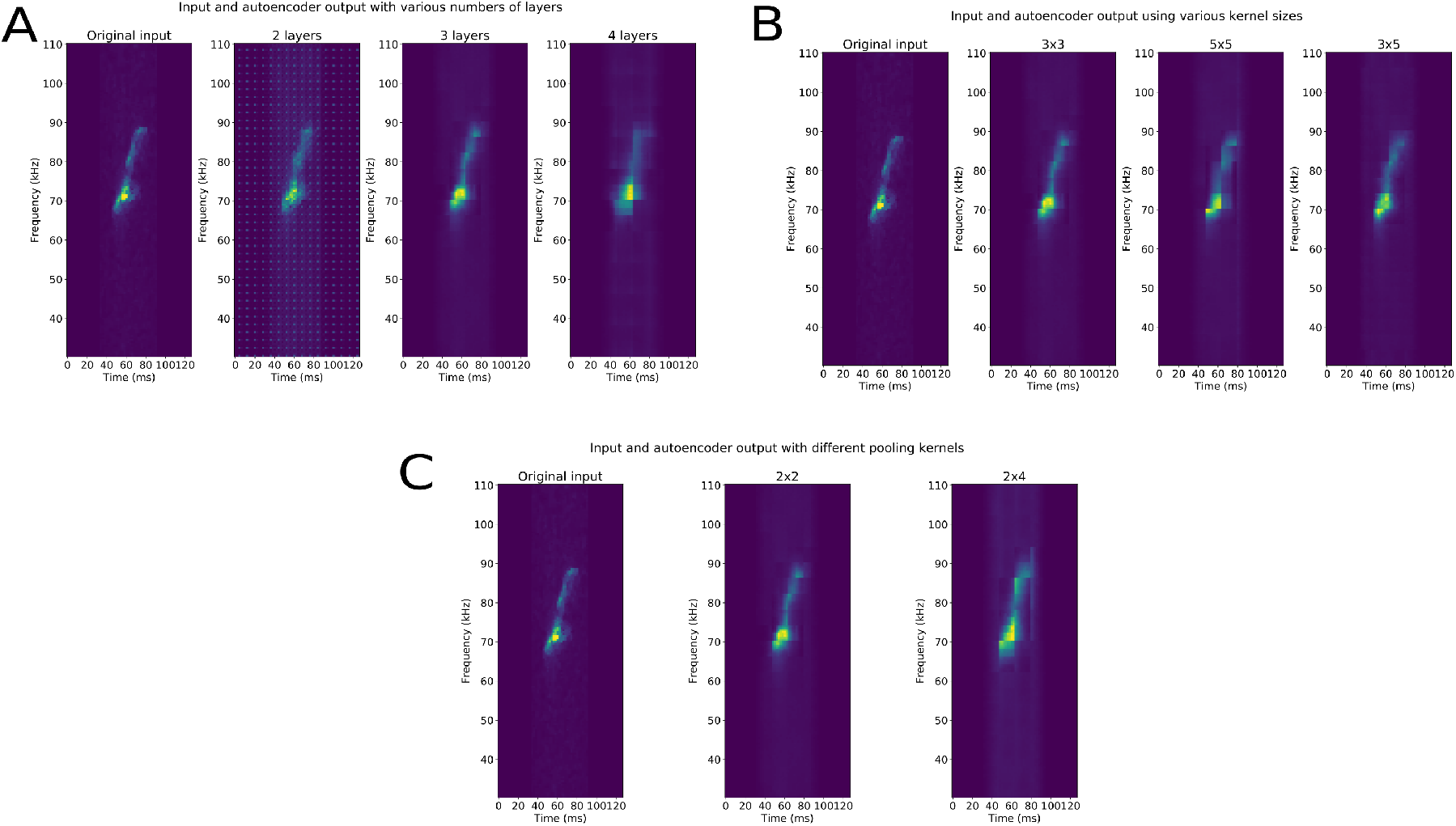
Effect of parameter tuning on the convolutional autoencoder reconstructions. Effect of changing A) different numbers of convolutional layers, B) kernel sizes, and C) pooling kernel sizes

